# Tracking gonadotropin-releasing hormone-mediated movement in-vivo using quantum dot nanoparticles: insights into pathways leading to sperm release

**DOI:** 10.1101/2024.10.07.617027

**Authors:** A.R. Julien, A.J. Kouba, D. Kabelik, J.M. Feugang, S.T. Willard, C.K. Kouba

## Abstract

As the number of conservation breeding programs for at-risk amphibians increase, application of assisted reproductive technologies (ARTs) has likewise grown, with hormone therapy being the most widely utilized technique. Significant interest has been raised for viable alternatives to traditional intracelomic injection for hormone treatment that may be simpler, require less hormone, induce a faster response and be more suitable for animals of different sizes or for minimal contact. We utilized quantum dot (QD) nanoparticles conjugated to GnRHa (GnRHa-QDs) as a means of monitoring exogenous GnRHa dispersal through living Fowler’s toads to investigate how delivery route may affect hormone pathway and efficacy after intracelomic, nasal, and oral administration. Real-time GnRHa-QD movement was tracked using in vivo imaging while hormone binding and QD aggregation were validated using fluorescence microscopy and measurements of spermiation responses following the administration of unconjugated GnRHa, unconjugated QDs, and GnRHa-QDs GnRHa alone was found to induce spermiation in 58.3%, 76.9% and 92.3% of males when administered by intracelomic, oral and nasal routes, respectively. When given through intracelomic injection GnRHa-QD caused spermiation and dispersed throughout the abdominal cavity to then aggregate within the brain, heart, liver, gastrointestinal (GI) tract, kidneys, and testes while unconjugated QDs did not. By contrast, GnRHa-QDs administered by oral or nasal routes were observed moving to the tympanic membrane and down the throat to the GI tract of the toads, but only nasal-administered GnRHa-QDs showed organ aggregation and were only found in the kidneys. Unconjugated QDs exhibited similar in vivo signal dispersal as their conjugated counterparts initially but did not aggregate in any organs over time nor induce spermiation. Through the use of novel QD hormone conjugates, in vivo imaging, and histology we were able to gain insight into how hormone pathways and efficacy might be impacted by delivery route which result in variations in physiological response to hormone treatment.

## 1. Introduction

In amphibian conservation breeding programs, exogenous hormone treatment is a primary mechanism for the elicitation of sexual behaviors and gamete production. The types of hormones used, and their routes of delivery, differ across programs and taxa. Delivery routes vary based on animal characteristics, hormone effect, and desired response time. For example, intravenous delivery is characterized by distribution throughout the body by the circulatory system, resulting in rapid action while conversely, subcutaneous injection has a relatively slow absorption rate (Jin et al. 2015). The predominant methods of hormone treatment used in amphibian assisted reproduction programs are intramuscular (caudata, Lampert et al. 2022), dorsal lymph sac, and intracelomic (also called intraperitoneal) injection (anura; Kouba et al. 2012b; Della Togna 2020; Shaidani et al. 2021). These routes of injection result in rapid absorption and physiological response due to the high vascularization in muscle tissue and the proximity of several major organs within the coelomic cavity (Jin et al. 2015; Mitchell 2009). Unfortunately, there is a sparsity of information regarding the pharmacokinetics of hormones once administered to live amphibians, nor are the mechanisms known through which these agents have systemic exposure or bioavailability. One reason for this lack of knowledge is due to challenges of following hormone movement in-vivo in live, whole animal systems.

While there are several studies that examine hormone binding kinetics and their physiological expression, few of these studies are performed in-vivo; rather, they frequently require the sacrifice of experimental animals and in-vitro tissue imaging. As a result, these studies lack insight into hormone pathways, timing, and efficacy, particularly regarding how these properties are impacted by administration method. Quantum dots (QDs) are a type of nanoparticle that provide a bright, long-lasting fluorescent or bioluminescent signal for detection within living system tissue, in contrast to many traditional fluorescent dyes and proteins. They are also versatile in function, application, color, and intensity (Probst et al. 2014) and when covalently linked to complex biomolecules such as peptides, proteins, and nucleic acids they are ideal for targeting specific tissues for in-vivo imaging (Xing and Rao, 2008; Gao et al., 2012). When conjugated to a hormone of interest and utilized with a positron emission tomography (PET)-scan like in-vivo imaging system (IVIS; Perkin Elmer), it may be possible to investigate hormone migration pathways and actions following traditional and experimental administration routes in an animal model.

Gonadotropin-releasing hormone analog (GnRHa) is commonly utilized in amphibian ART programs worldwide (Browne et al. 2006; Kouba et al. 2009; Clulow et al. 2018; Silla et al. 2019). In the US, for example, its use is standard practice in breeding programs for species such as the federally listed Puerto Rican crested toad (Burger et al. 2021) and the dusky gopher frog (Kouba et al. 2012a; Graham et al. 2018), among many others. Both endogenous and exogenous GnRH are used to stimulate a natural cascade of gonadotropins in amphibians by directly activating the hypothalamic-gonadal-pituitary (HPG) axis (Daniels and Licht 1980; Vu and Trudeau 2016), leading to the increased production and expulsion of sperm in males and follicular development and ovulation in females (Licht et al. 1987; Browne et al. 2006; Brown and Zippel 2007; Kouba et al. 2009; Graham et al. 2018; Bronson et al. 2020). Because of its prolific use, GnRHa has seen the greatest experimentation in alternative routes. In addition to the traditional intramuscular and intracelomic injection routes, dermal (Silla et al. 2020; Campbell et al. 2021), oral (Rowson et al. 2001; Chen et al. 2024), and nasal (Julien et al. 2019) delivery routes of GnRHa have been tested in a small number of both anuran and caudate amphibians. Responses to GnRHa given through these delivery methods has been variable across studies, likely due to substance pathway migration; isozymes of GnRH receptors are distributed across numerous tissues in amphibians with the primary high-affinity receptors for GnRH localized within the anterior pituitary, providing many possible locations of action for exogenous GnRH. Thus, the route of administration of exogenous GnRHa may play a role in how quickly, and at what concentration, bioavailable hormone reaches receptors and causes downstream effects.

The use of in vivo tracking with hormone-conjugated QDs may provide an opportunity to build understanding of the pathways and tissues through which hormones such as GnRHa are transmitted following alternative routes of administration, particularly in comparison to the traditional intracelomic route. The large volume of literature regarding the use of GnRHa for eliciting gamete release makes it an ideal conjugate for in vivo QD monitoring by providing an additional physiological validation of hormone binding following administration; while visualization of QD fluorescent signal will provide information regarding the pathways intracelomic, oral, and nasal delivery follow, subsequent spermiation responses will indicate hormone binding and receptor activation in the areas where QD signal is also found. Here we used GnRHa-conjugated quantum dots (GnRHa-QD) for the first time to map in-vivo GnRHa migration pathways and organ targeting to gain information on reproductive hormone efficacy in an amphibian. Our goals were to: a) visualize pathways following intracelomic, oral, and nasal delivery routes using GnRHa-conjugated fluorescent quantum dots with in vivo imaging and histology, and b) compare efficacy of GnRHa between these routes and validate imaging pathways by measuring spermiation response.

## 2. Materials and Methods

### 2.1 Study Animals

Sexually mature male Fowler’s toads (*Anaxyrus fowleri*) were collected from Oktibbeha County in Mississippi during the breeding season from April-July 2016 and 2017 (Permit #0728161), March-April 2018 (Permit #0501181), and April 2021 (Permit #0504202). Males weighed on average 24.0 ± 0.8 grams. Toads were housed in accordance with the Institutional Animal Care and Use Committee (IACUC) protocols at Mississippi State University (IACUC #16-406; #19-345). Briefly, males were housed in polycarbonate tanks (30 cm L x 46 cm W x 66 cm H) in groups of 3-6 on a substrate of cocoa fiber and soil cultured with springtails (Collembola), with terra cotta pots and PVC piping as shelter options. Animals were fed a diverse diet of crickets, Dubia roaches, and mealworms three times a week on a rotating schedule. Prior to feeding, all insects were gut-loaded with vitamin supplements (Repashy Ventures Inc., CA, USA) and coated in Reptivite© calcium powder (ZooMed Laboratories Inc., Costa Mesa, CA, USA). Water was available ad libitum. Animal housing areas were kept on a 12-hour day/night cycle and were maintained at 23 ± 2 °C.

### 2.2 Spermiation Trials Trial 1

#### Trial 1

To validate the efficacy of GnRHa at eliciting spermiation through varying routes, male toads were treated with a single 10 µg dose of GnRHa (Sigma-Aldrich #L4513) that was administered either intracelomically (n = 12), intranasally (n = 13), or orally (n = 13) (Figure 1). GnRHa was dissolved in a vehicle of 200 *µ*l phosphate buffered saline (PBS) and delivered by intracelomic injection into the coelom of male toads using a 30-gauge insulin needle and syringe. For intranasal and oral delivery, GnRHa in a vehicle of 10 *µ*l of PBS was pipetted directly into the nares or mouth of individuals while they were held upright. Toads were held in this position for approximately 5 seconds following treatment to allow hormones to drain down the nares or throat.

**Figure 1.**
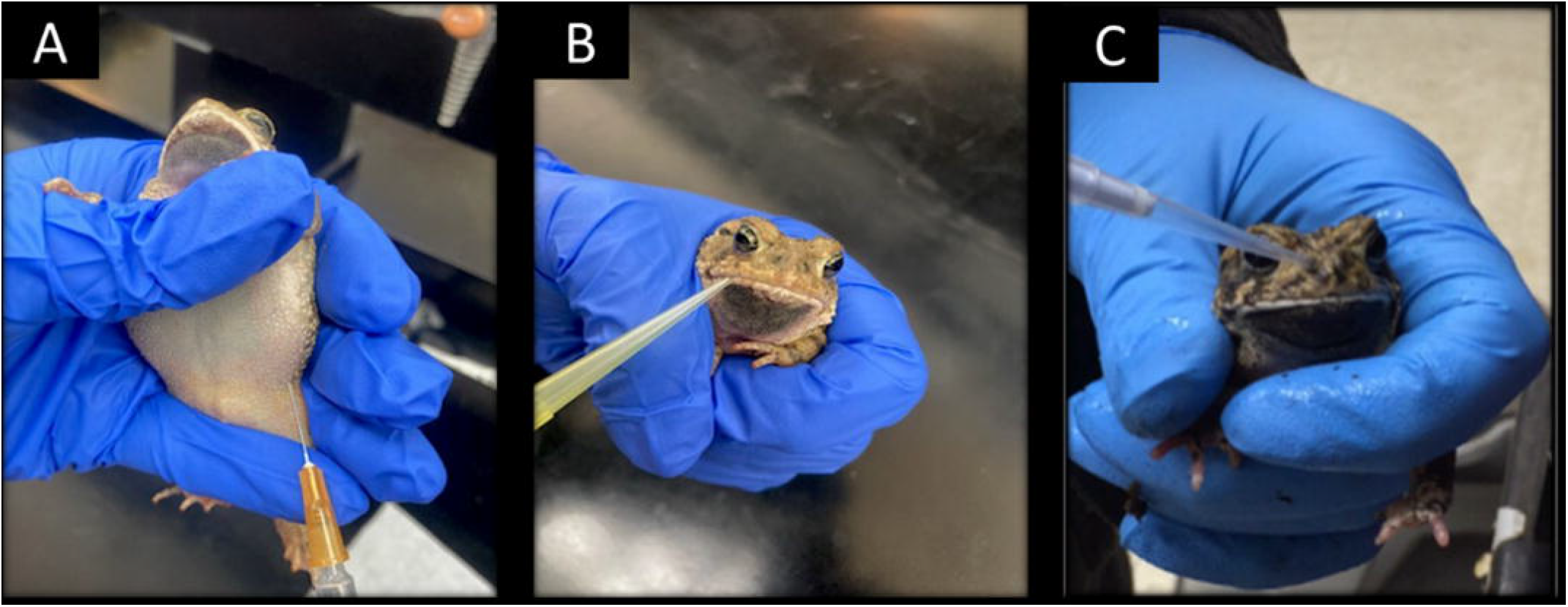
Animal handling methods during intracelomic (A), oral (B), and nasal (C) hormone delivery

#### Trial 2

A second spermiation test was conducted to determine the ability of QDs to elicit a spermiation response. Additional males (n = 18) were administered either 50 nM of unconjugated quantum dots (QTracker 655 Fisher Scientific/Invitrogen; Q25021MP, excitation/emission: 455-585/655, 15-20 nm) or GnRHa-conjugated quantum dots (GnRHa-QD; GnRHa-QTracker 655 Fisher Scientific/Invitrogen) via intranasal (n = 3 males/QD treatment), intracelomic (n = 3/trt), or oral (n=3/trt) administration routes, and assessed for sperm release and quality. Exact ratios of GnRHa to QD conjugations were not made available from Invitrogen, but were estimated at 0.17 µg GnRHa per 0.4 nM of QDs. Delivery methods were the same as described above.

### 2.3 Collection and Analysis of Spermic Urine

Following treatment with GnRHa, unconjugated QDs, and GnRHa-QDs, we compared the number of males producing sperm, sperm quantity, and sperm quality of each spermic urine sample between the three different routes of hormone delivery over time. Urine was collected by holding males upright over sterile petri dishes and urination occurred through a natural response to a perceived threat. Samples were collected hourly for eight hours to determine the latency of sperm production between treatment routes. Urine volume was measured at each time point and analyzed using an Olympus CX-41 phase contrast microscope for sperm presence, motility, and concentration. Sperm motility was quantified using a multi-cell counter. Sperm were categorized by type of movement exhibited: forward progressive motile (FPM) were sperm exhibiting flagellar movement propelling them forward; motile (M) sperm exhibiting flagellar movement but remaining stationary; and non-motile (NM) sperm which were stationary with no flagellar movement. Forward progressive, motile, and non-motile sperm were counted until 100 cells were reached and the ratio of each category were recorded as percentages. To calculate total motility (TM), we added FPM and M sperm together. Sperm concentration was evaluated using a hemocytometer (Hausser Scientific #3200), wherein all sperm cells within specific grid regions were counted using a cell counter.

### 2.4 IVIS Imaging

A subset of males treated with unconjugated QDs (n = 2/route) and GnRHa-QDs (n = 2/route) from the second spermiation trial were utilized for live-animal imaging. To visualize fluorescence signal from QDs within the live animal over time, males were imaged using an in-vivo imaging system. Prior to QD administration and imaging, male toads were anesthetized using ethyl 3-aminobenzoate methanesulfonate (MS-222; Sigma-Aldrich E10521) at a concentration of 1 g/L for 30 minutes. Anesthetized toads were placed into the In-Vivo-Imaging System (IVIS; Perkin-Elmer) and fluorescence detection was performed using excitation filters of 540, 560, 580, 600, and 620 nm. Toads were imaged both dorsally and ventrally. Toads were imaged prior to QD-conjugated hormone administration as a control and at 1- and 24-hours post-treatment. Once fluorescent images were obtained by the IVIS, spectral unmixing was performed with the Living Image software (Perkin-Elmer) in order to select the most accurate detection filter (570 nm).

### 2.5 Euthanasia and Histology

Further investigation of QD pathways was conducted through visualization of organ aggregation. Following IVIS imaging, males were euthanized at either the 1-hour (n =1 male /route of administration) or 24 hour (n=1/route) time points post-GnRHa-QD administration using an overdose of MS-222 at a concentration of 3 g/L for 60 minutes. The brain, heart, liver, a section of GI tract, kidneys, and testes were excised for comparison of tissue fluorescence signal accumulation and localization following nasal, oral, and intracelomic GnRHa-QD delivery. Briefly, histological samples were encased in paraffin wax and sectioned at a thickness of 5 *µ*m, which were then mounted onto slides. Slides were deparaffinized and mounted using DAPI fluorescent mounting media (VECTASHIELD; excitation/emission: 360 nm/ 460 nm) prior to cover slipping. Subsequently, tissue sections on slides were imaged for fluorescence signal using a Nikon 80i microscope with images assessed at 20x magnification and a QD-specific filter cube (485 - 525 nm excitation) plus a DAPI filter cube (352 - 402 nm excitation).

### 2.6 Statistical Analysis

Sperm production was compared between the three delivery methods, intracelomic, intranasal, and oral administration. Assessed spermiation parameters included sperm concentration (sperm/mL), forward progressive sperm motility (FPM), and total sperm motility (TM). Analyses of parameters included male ID and collection time point as random effects; however, these effects were removed as they did not explain any additional variation in the models. Final analyses for all parameters were run using a generalized linear model (GLM), with administration type (intracelomic, intranasal, and oral) as a fixed effect. FPM and concentration were run under a gamma family with an inverse link, and TM was run under a gaussian family with a log link. If significance was found in the models, a post-hoc analysis using the estimated marginal means (emmeans package within R) was run to determine specific differences between the hormone administration types. We also compared the percentage of responders separated by administration route to determine if there was any variation in overall response success between routes. We ran a Generalized Additive Model for Location Scale and Shape (GAMLSS) using a beta zero-inflated family, with administration routes (oral, intracelomic, and intranasal) and time as fixed effects. We assessed normality and homogeneity of variance using Shapiro-Wilke’s and Levene’s test prior to running models. Statistical significance was set at 0.05, and all data were analyzed in RStudio.

## 3. Results

### 3.1 Effects of GnRHa administration route on sperm quantity and quality

Following GnRHa administration, spermiation was observed at the first time point, 1-hour post-treatment following each delivery route. Analysis revealed a treatment effect, with intranasal delivery of GnRHa resulting in a significantly higher (92.3%; p < 0.05) percentage of responding males than both oral delivery (76.9%; t = - 4.072) and IP injection (58.3%; t = - 6.420). Furthermore, significantly more males responded to oral delivery (p = 0.019; t = 2.520) than intracelomic injection. The highest proportion of males responded to intranasal delivery at 1-hour post-treatment, while the highest percentage of males responded to oral delivery at 4 and 5 hours. The highest average proportion of responders following intracelomic injection did not occur until 8 hours post-treatment (Figure 2). No significant effects were observed as a result of time, nor was there a time x treatment interaction effect (p > 0.05). While not significant, trends were observed in latency to spermiation; intracelomic injection had the highest average latency compared to oral or intranasal administration (3.1 hours vs. 1.9 and 1.8 hours, respectively). Thus, intranasal administration resulted in the fastest spermiation response and had the highest percentage of responding males, while intracelomic injection resulted in the lowest percentage of males producing sperm and they responded at a much later point in time (Table 1).

**Table 1:**
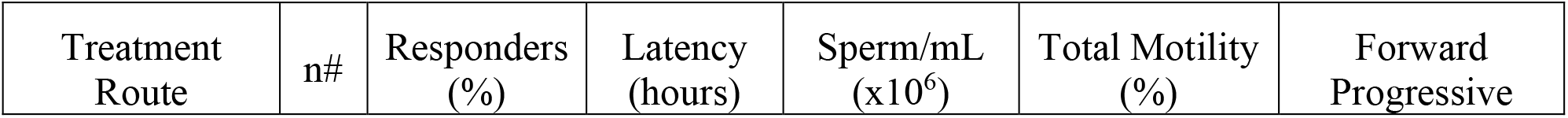
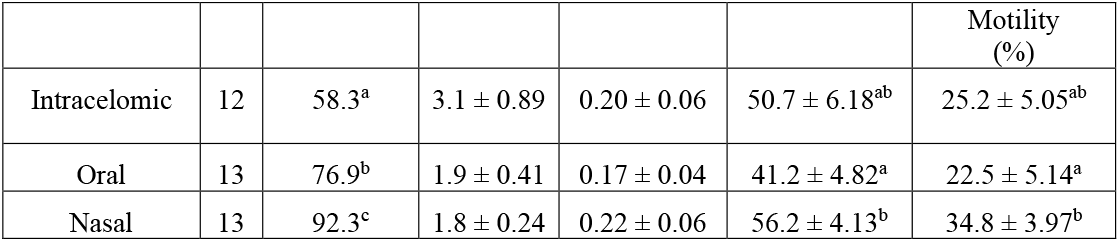
Average number of responders and sperm parameters following intracelomic, oral, and nasal administration of 10 µg GnRHa. Data are shown as mean ± SEM, and superscripts denote significant differences.

| Treatment Route | n# | Responders (%) | Latency (hours) | Sperm/mL (x10<sup>6</sup>) | Total Motility (%) | Forward Progressive |
| --- | --- | --- | --- | --- | --- | --- |

|  |  |  |  |  |  | Motility<br>(%) |
| --- | --- | --- | --- | --- | --- | --- |
| Intracelomic | 12 | 58.3<sup>a</sup> | 3.1 ± 0.89 | 0.20 ± 0.06 | 50.7 ± 6.18<sup>ab</sup> | 25.2 ± 5.05<sup>ab</sup> |
| Oral | 13 | 76.9<sup>b</sup> | 1.9 ± 0.41 | 0.17 ± 0.04 | 41.2 ± 4.82<sup>a</sup> | 22.5 ± 5.14<sup>a</sup> |
| Nasal | 13 | 92.3<sup>c</sup> | 1.8 ± 0.24 | 0.22 ± 0.06 | 56.2 ± 4.13<sup>b</sup> | 34.8 ± 3.97<sup>b</sup> |

**Figure 2.**
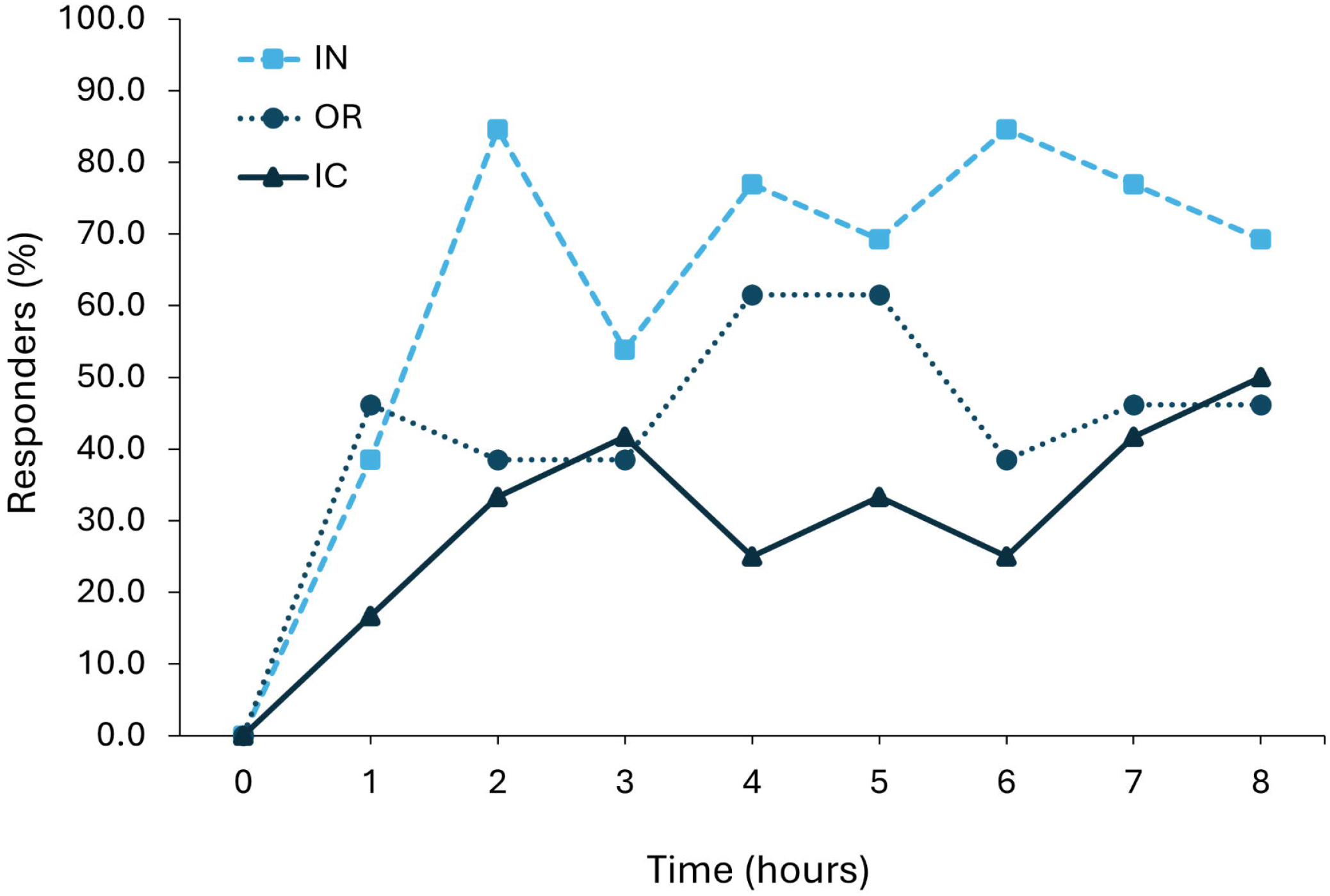
Percentage of males producing sperm in response to 10 µg GnRHa via intracelomic (IC; n = 12; ▲), oral (OR; n = 13; ●), and intranasal (IN; n = 13; ■) routes over time.

Hormone delivery route did not have a significant (p > 0.05) effect on the measured concentration of sperm/mL across groups. (Table 1). While high percentages of sperm motility were observed soon after hormone treatment, across all hormone delivery routes, concentration of sperm/mL did not peak until several hours post-treatment. Though not significant, concentration of sperm/mL is highest at 5-hours post-treatment for both intranasal (0.50 × 106) and oral (0.39 × 106), while the highest concentration of sperm following intracelomic injection (0.59 × 106) was not reached until time point 7 (Figure 3).

**Figure 3.**
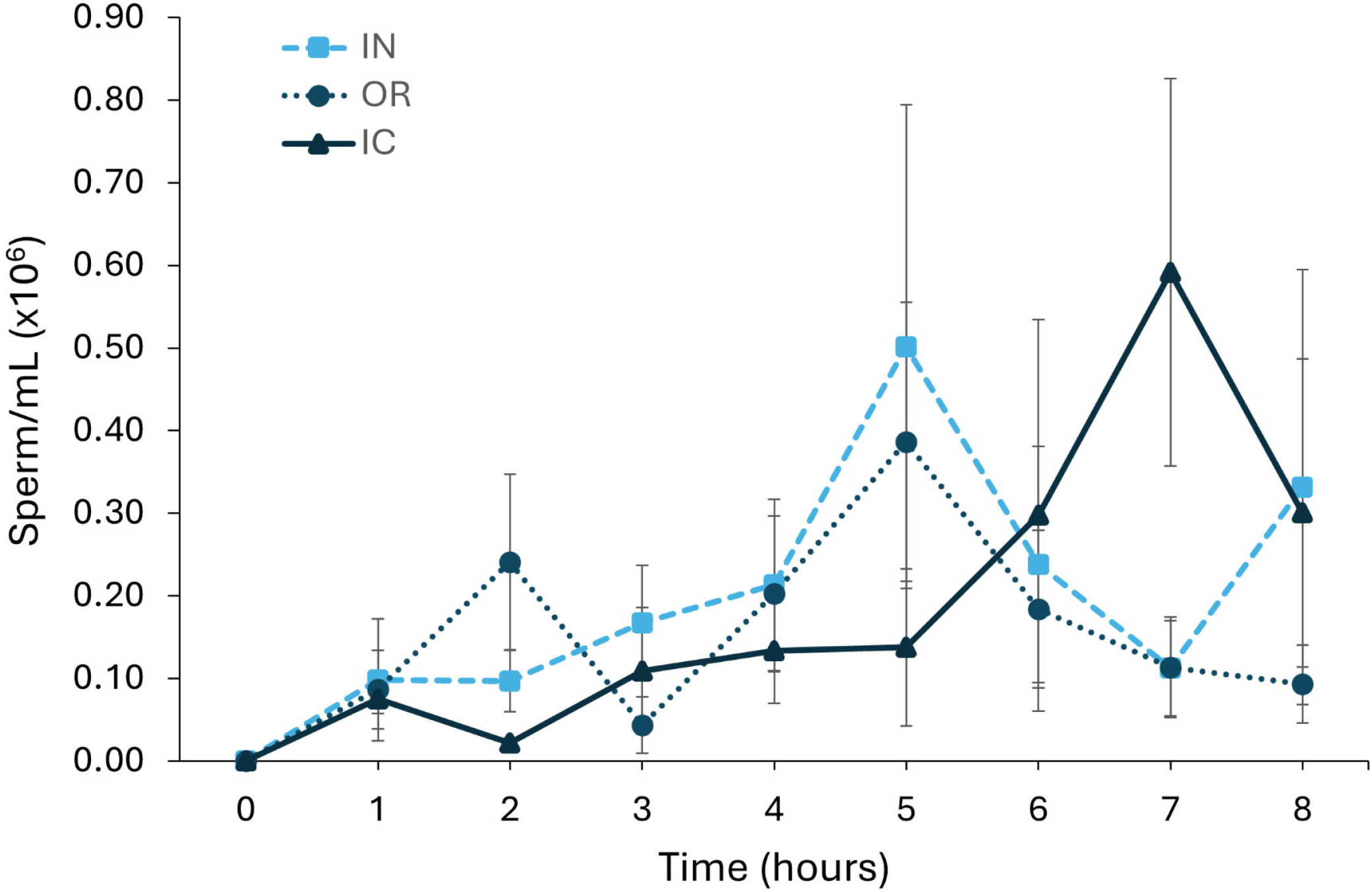
Average concentration sperm released over time for male toads administered 10 µg GnRHa via intracelomic (IC; n = 12; ▲), oral (OR; n = 13; ●), and intranasal (IN; n = 13; ■) routes over time. Data are shown as mean + SEM.

The highest percentage TM sperm following intracelomic injection occurred by the first hour after treatment (Figure 4a), while the highest percentage of FPM sperm occurred at the same time the highest sperm concentration, 7-hours post-treatment (Figure 4b). Percentage of FPM and TM sperm were similar (FPM: p = 0.125; TM: p = 0.455) whether GnRHa was delivered intracelomic or nasally. However, nasal delivery resulted in significantly higher (t = 2.491; p = 0.014) percentages of FPM sperm and higher (t = -3.473; p = 0.001) percentages TM sperm than oral administration. Intracelomic delivery of GnRHa did not significantly differ from oral administration in either sperm FPM (p = 0.831) or TM (p = 0.108). While time was not a significant factor, fluctuations in sperm motility over time were observed, as with sperm concentration. Following intranasal delivery of GnRHa, sperm FPM (46.5%) and TM (74.9%) were highest at 4 and 5-hours, respectively. In contrast, sperm TM (67.5%) was highest at 3-hours for oral delivery, while the highest sperm FPM (29.3%) was not observed until 8 hours following hormone administration.

**Figure 4.**
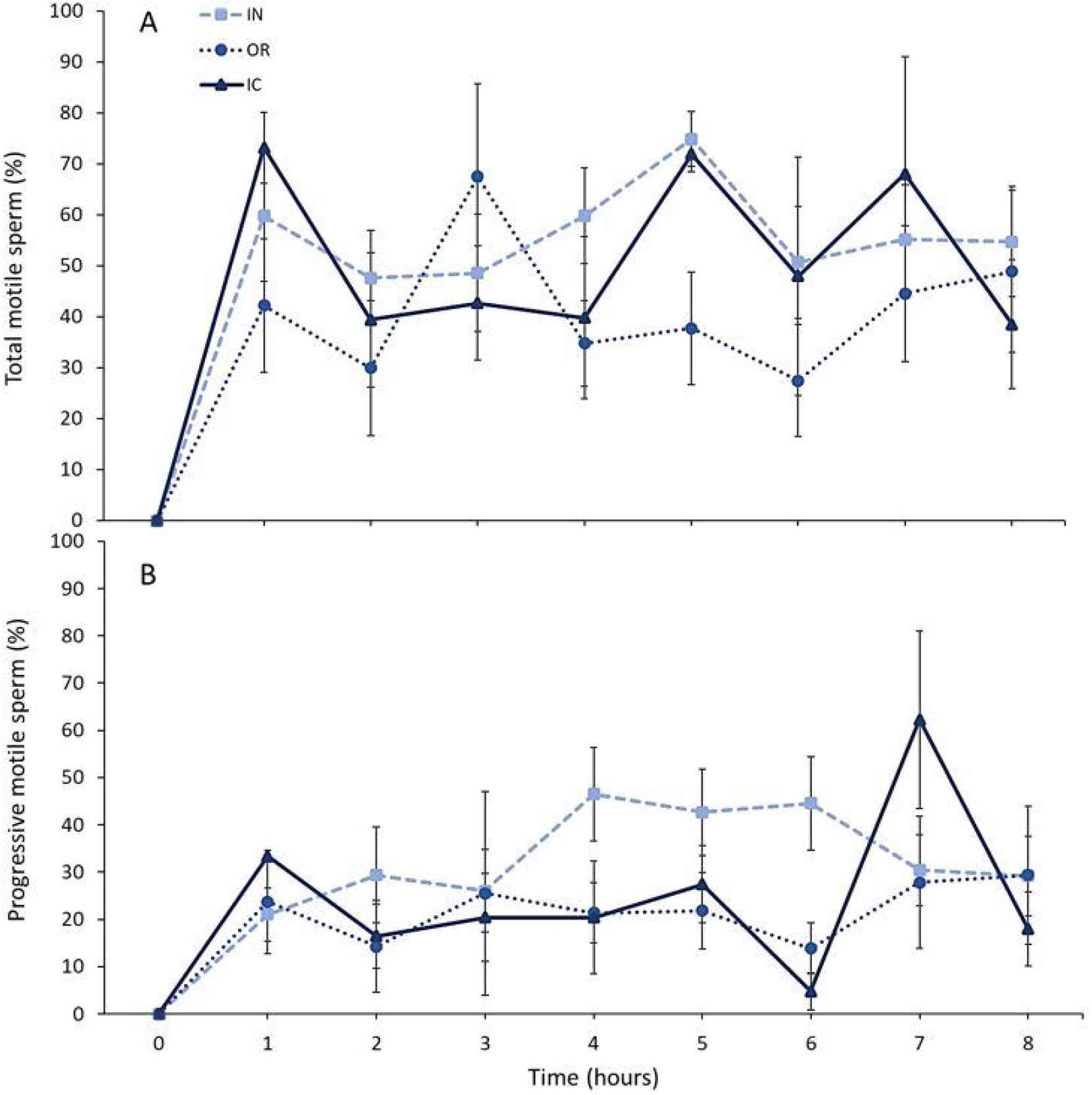
Forward progressive motility (A), and total motility (B) of sperm samples collected from male toads given 10 µg GnRHa via intracelomic (IC; n = 12; ▲), oral (OR; n = 13; ●), intranasal (IN; n = 13; ■) routes over time. Data are shown as mean + SEM.

Intracelomic injection of GnRHa-QDs induced spermiation in 100% of males, with an average sperm concentration of 3.84 × 104 ± 1.21 × 104 sperm/mL and average motility of 47 ± 7 %, while unconjugated QD did not induce any spermiation response. Nasal and oral administration of either unconjugated or GnRHa-QDs resulted in no detectable sperm release from the three animals tested.

### 3.2 IVIS Imaging of quantum dots

In-vivo imaging revealed substantial visible QD fluorescence signal (displayed here with a synthetic color scale of yellow to red, with yellow indicating the highest aggregation of QDs and red the lowest) within GnRHa-QD treated toads. The male given GnRHa-QDs via intracelomic injection showed fluorescence signal in the lower back region 1-hour after treatment when imaged dorsally, and on near the site of injection when imaged ventrally (Figure 5a). In vivo imaging of the dorsal side of males 1-hour following GnRHa-QD treatment showed fluorescence aggregation at the tympanum from both oral (Figure 5b) and intranasal delivery (Figure 5c), with additional fluorescence signal observed at the nares of the intranasal-treated male. In addition, ventral imaging revealed fluorescence signal at the mouth of both oral and intranasal treated males, with oral delivery exhibiting signal at the throat.

**Figure 5.**
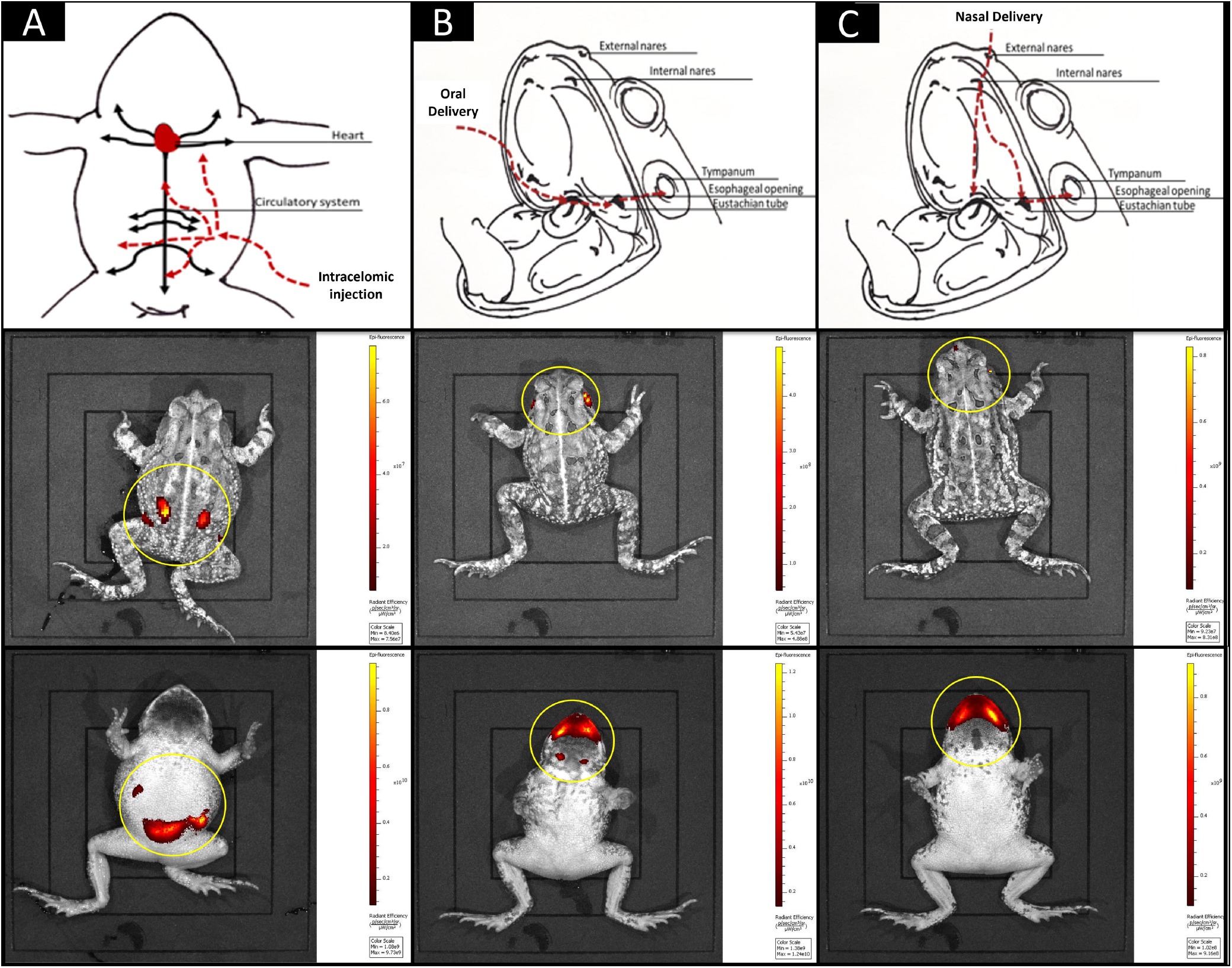
Proposed exogenous hormone pathways and visible GnRHa-QD fluorescent signal at 1-hour following intracelomic (A), oral (B), intranasal (C) delivery. Yellow circles indicate GnRHa-QD localization.

Movement and localization of hormone conjugated nanoparticles was very different between the 1- and 24-hour time points with in-vivo imaging of fluorescent signal following administration of GnRHa-QDs showing clear differences in nanoparticle dispersal depending on delivery route. Intracelomic injection predictably resulted in localized observed fluorescence at the point of injection in the abdomen at 1-hour post-delivery (Figure 6a), with GnRHa-QD signal dispersing across the abdomen by 24 hours post-treatment (Figure 6d). However, fluorescence was observed at the mouth and throat at 1-hour post-oral delivery (Figure 6b) in ventral images but only observed at the lower abdomen and gastrointestinal (GI) tract within 24 hours (Figure 6e). Similarly, nasal delivery of GnRHa-QDs presented with observable fluorescence at the mouth directly below the internal nares at 1-hour post-treatment (Figure 6c), with signal migrating to the lower abdomen and GI tract at 24 hours post-treatment (Figure 6f).

**Figure 6.**
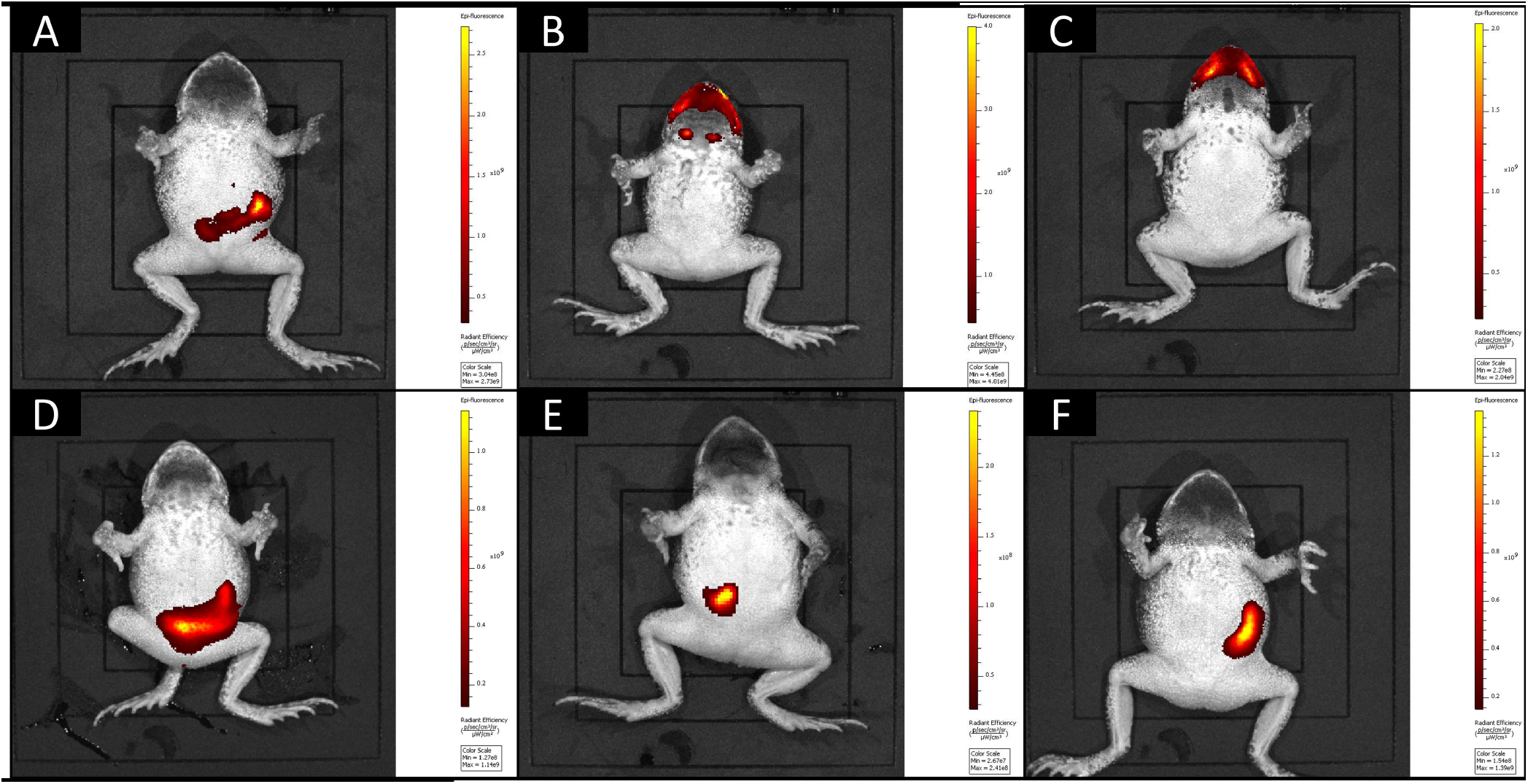
Visible GnRHa-QD fluorescence at 1-hour (top row) and 24-hours (bottom row) following intracelomic (A & D), oral (B & E), and intranasal (C & F) delivery.

Unconjugated QD signal showed similar migration patterns as GnRHa-QD signal across administration routes with the exception that dorsally, unconjugated QDs were not detected at either 1 hour or 24 hours post-treatment (images not shown). No fluorescence signal was detected in males imaged at baseline prior to QD administration.

### 3.3 Histology

Histological analysis revealed differences in localization of GnRHa-QD aggregation between treatment routes and within tissues and organs (Figure 7). Only intracelomic injection resulted in GnRHa-QD aggregation in all excised organs including: the brain (Figure 7a), heart (Figure 7b), liver (Figure 7c), GI tract (Figure 7d), kidney (Figure 7e), and testes (Figure 7f). Localization of GnRHa-QDs was observed in all organs at 1-hour post-intracelomic injection, whereas by 24-hour, fluorescence was only observed within the kidneys (Figure 8a-b). Unconjugated QD fluorescent signal was detected only in the kidney following IP injection at hour 24. In males treated intranasally with GnRHa-QDs, signal was detected only within the GI tract, and was observed at both 1-hour and 24-hours post-treatment (Figure 8c-d). Intranasal administration of unconjugated QDs showed no fluorescent signal at any of the imaging time points. These results stand in stark contrast with the oral delivery of hormone, where no GnRHa-QD signal as observed in any of the excised organs selected or histology at either the 1-hour or 24-hour time points.

**Figure 7.**
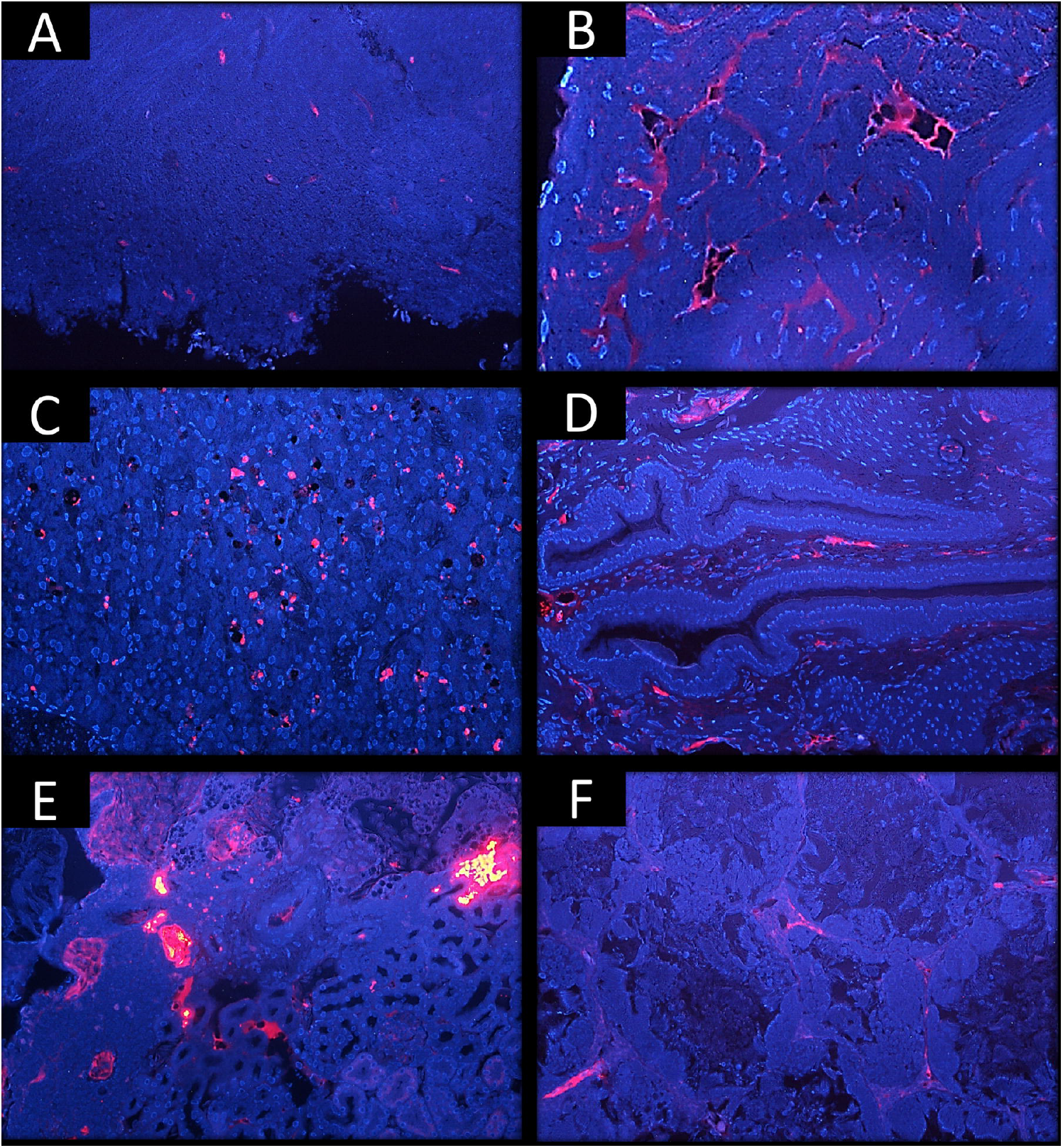
Aggregation of GnRHa-QDs in histological sections of the brain (A), heart (B), liver (C), GI-tract (D), kidney (E), and testis (F) one-hour following intracelomic injection. DAPI DNA staining shown in blue. GnRHa-QD aggregation shown in red.

**Figure 8.**
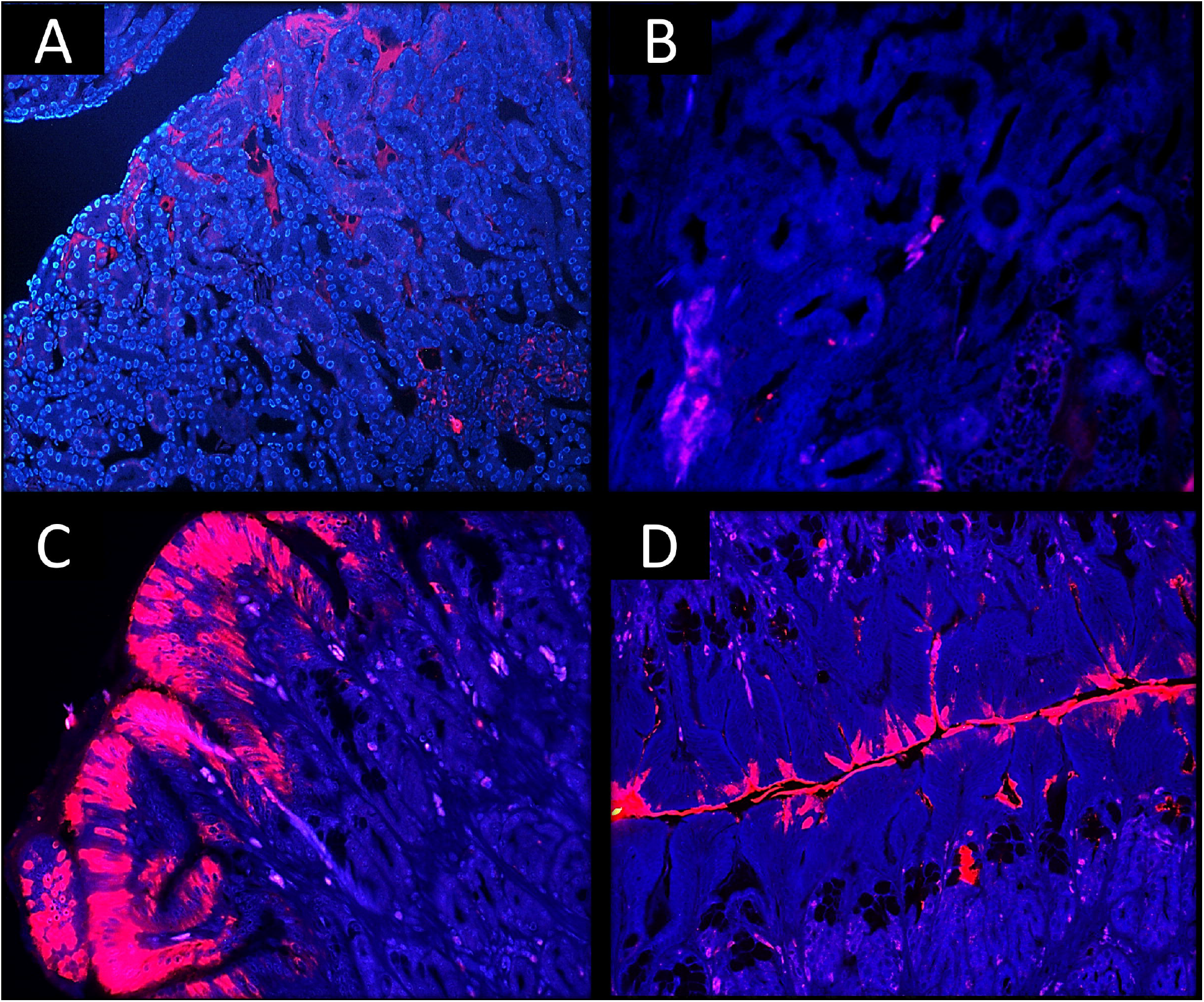
Histological sectioning of toad kidney following intracelomic injection of GnRHa -QDs at (A) 1 hour, and (B) 24 hours post-treatment and histological sections of the GI tract following intranasal administration at (C) 1 hour and (D) 24 hours post-treatment. DAPI DNA staining shown in blue. GnRHa-QD aggregation shown in red.

## 4. Discussion

Intracelomic injection of GnRHa and GnRHa-QDs provided results consistent with known intracelomic pathways that lead to spermiation, but also demonstrate spatial and temporal distribution of GnRHa. Following injection into the coelom, exogenous substances such as GnRHa and GnRHa-QDs are absorbed into an expansive capillary network throughout the body cavity as the coelom has a large surface area and is highly vascularized. Additionally, several organs of interest in this study reside directly within the coelomic cavity including the heart, liver, intestinal tract, kidneys, and testes. As a result, substances are capable of circulating through the bloodstream and directly reaching various organs and sites of action (Wright and Whitaker 2001; Helmer and Whiteside 2005; Vieu et al. 2024). Imaging of live toads revealed a clear dispersal of GnRHa-QD fluorescence signal throughout the body cavity immediately following intracelomic delivery, and histological analysis showed GnRHa-QD signal in all examined organs (brain, heart, liver, GI tract, kidney, and testes) in areas with dense vascularization after just 1 hour (Figure 7). After 24 hours, GnRHa-QDs were only found in the kidneys (Figure 8). Altogether, it appears that GnRHa administrated at the coelom enters the bloodstream to rapidly disperse through numerous organs and is likely filtered through the kidneys for excretion within 24 hours.

Intranasal and oral administration of GnRHa also resulted in spermiation from A. fowleri, at more reliable rates than the traditional intracelomic approach. While nasal administration of GnRHa was previously found to be an effective means of eliciting spermiation in amphibians (Julien et al. 2019), spermiation results following oral administration have been variable (Rowson et al. 2001; Chen et al. 2024). While sperm concentration was similar across delivery routes, sperm motility (forward progressive and total sperm motility) and overall male response were best when GnRHa was delivered intranasally. Live imaging of GnRHa-QDs between these routes demonstrated similar migration patterns. Immediately post-treatment, both pathways exhibited fluorescence signal from GnRHa-QDs at the tympanum and mouth, with nasal delivery showing additional signal at the nares. These migration patterns align with known anuran physiology. Anurans possess internal nares that open into the mouth; thus, substances delivered at the external nares can drain through the internal nares and into the mouth, where they may subsequently flow through the eustachian tubes to both the esophagus and tympanum. This is further supported by visible fluorescence detected within the abdomen at 24-hours post-treatment, in areas consistent with gastrointestinal (GI) tract location (Figure 5d-e). Interestingly, fluorescence was detected within the GI tract upon histological examination in males given GnRHa-QDs through the nares, but not following oral administration; fluorescence from unconjugated QDs was not observed in the GI tract after either nasal or oral administration. A lack of observed signal from GnRHa-QDs following oral administration may be because swallowed substances travel to the GI tract for absorption by the stomach and intestines. While the GI tract has a large, highly absorptive surface area, bioavailability of administered substances can be hindered by digestive enzymes and low pH of the gastric environment and the various cellular layers that substances must pass through (Homayun et al. 2019). It should also be noted that live-imaging revealed fluorescence signal from nasal and oral administration at opposite sides of the abdomen; this may indicate that these two routes were absorbed at different rates over the 24-hour time course and reached different areas of the GI tract. Finally, only small segments of the GI tract were taken for histological examination, and it is possible that the segments wherein orally administered GnRHa-QDs were absorbed were not selected. Overall, live imaging and histological examination following nasally and orally administered GnRHa and QDs reveal that these substances are capable of reaching multiple tissues following delivery within 24 hours before reaching the GI tract.

While the intracelomic pathways revealed through imaging and histology were consistent with the literature surrounding intracelomic drug administration, the physiological response following administration of GnRHa via this route resulted in the lowest percentage of responding males and a higher latency to sperm parameter peaks. In contrast, both nasal and oral delivery routes resulted in more male responders and higher sperm concentration earlier than intracelomic injection. These unexpected lower responses may be due to the time needed for intracelomic injected GnRHa to reach the pituitary, and possible reduction in GnRHa concentration following dissipation through coelom and into the bloodstream. The primary binding site for GnRH is the pituitary; however, studies have noted additional receptors within the brain and the gonads (Millar 2003; Millar 2004; Ramakrishnappa et al 2005; Anjum et al. 2013). Additionally, GnRHa may exert its action on the pituitary following diffusion into the bloodstream regardless of administration route, but the presence of the gonads within the coelom should have resulted in hormone binding and rapid response if GnRHa receptors were available on the gonads. However, because both nasal and oral routes resulted in a faster spermiation response, coupled with a lack of GnRHa-QD signal within the gonads, it is likely that GnRHa given through intracelomic, oral, and nasal administration all targeted the receptors on the pituitary, and when GnRHa is administered through oral or nasal routes that the proximity of mucosal absorption at the nares, mouth, and throat to the bloodstream and pituitary allowed for an earlier spermiation response. Due to the greater dilution of GnRHa through the coelomic cavity and into the bloodstream necessitated by intracelomic injection, it is likely that a lower concentration of GnRHa ultimately reached the pituitary compared to orally and nasally administered hormone. If exogenous hormones are able to cross into the blood stream near the oral or nasal cavities, then the distance of hormone travel would be reduced and the concentration of hormone reaching the pituitary may be increased.

Interestingly, intracelomic injection of GnRHa-QDs was the only route to result in an observed spermiation response. Because unconjugated QDs themselves did not elicit a sperm reaction following any of the treatment routes, but intracelomic injection of GnRHa-QDs did, some concentration of bioavailable GnRHa was able to bind to receptors while conjugated to QDs. This is possibly due to a vascular route taken by intracelomic injection compared to the routes taken by oral and nasal administration. Intracelomic injection, as previously discussed, is an administration route that allows for large concentrations of bioavailable hormones and pharmaceuticals to be absorbed into the bloodstream and surrounding organs due to the large surface of the coelomic cavity and its extensive vascular network. It is also possible that the GnRHa-QDs administered into the coelom were not able to reach the pituitary and instead bound to receptors on the gonads. This may also explain the comparatively weak spermiation response that resulted from intracelomically-administered GnRHa-QDs compared to GnRHa alone.

By contrast, oral administration must pass through both the GI tract and the liver before dispersal through the bloodstream, thus decreasing the overall concentration of GnRHa-QDs that reach the bloodstream by breaking the conjugate down in the GI tract or filtration by the liver. Although intranasal administration is also known to act in a more localized fashion, having a reduced systemic effect (Homayun et al. 2019), IVIS imaging revealed that intranasal administration, similar to oral administration, was largely swallowed. One of the drawbacks in medical applications of oral delivery is that absorption through the GI tract is a relatively slow process with the potential of breaking down administered compounds (Al Shoyaib et al. 2020). Thus, the breakdown of GnRHa-QDs by the GI tract may account for a lack in spermiation response from both nasal and oral routes during sperm collection points.

As a result of the conjugation process, the exact concentration of GnRHa within QD solutions were unable to be determined; it is possible that the conjugation process reduced the concentration of bioavailable GnRHa or the resulting conjugate was larger than both GnRHa of QDs alone. While the vascular route taken by intracelomic injection may allow for larger substances, as well as larger concentration of substances, to reach the pituitary via the bloodstream, the routes taken by oral administration and nasal administration may be more restrictive in size. Because intranasal administration of GnRHa alone resulted in a higher proportion of responding males compared to oral administration, it is likely that a sufficient concentration of the hormone is transported through the vascularized mucosal layer of the nares to enter the bloodstream near the pituitary, in addition to being swallowed, which may account for the higher response than that of oral administration. However, a lack of spermiation following intranasal administration of the GnRHa-QD conjugates may indicate that an insufficient concentration of GnRHa is able to reach the targeted receptors through this method, either because the conjugates were too large to cross the mucosal layer, or that the concentration of GnRHa in the conjugates was insufficient to generate a response. It should also be noted that IVIS imaging revealed that both intranasal and oral administration of GnRHa-QDs dispersed into multiple locations following administration. In both routes, visible fluorescence was observed at the tympanum, mouth, and throat, with additional signal observed at the nares following nasal administration. Signal was observed at these locations nearly simultaneously, indicating that there was a large dispersal of GnRHa-QDs to multiple locations, and thus, areas of potential absorption. This may have further reduced the concentration of bioavailable GnRHa that was absorbed into circulation at each location. The overall size of the conjugates, coupled with the loss of significant volumes of administered substance to the GI tract and other locations following oral and nasal routes, may have caused too little bioavailable GnRHa to reach the pituitary compared to the volume that entered the bloodstream via intracelomic injection. However, because unconjugated QDs did not result in spermiation following any of the tested delivery routes nor were they found upon histological examination, we conclude that GnRHa-QD conjugations may have resulted in organ-specific aggregation despite a lack of physiological response.

Here, we present the first use of quantum dot nanoparticles for the study of hormone pathways in an amphibian model. Our results validate the use of QDs for investigating hormone pathways within a living system by providing evidence that administered GnRHa may follow separate routes through the body, but ultimately likely binds to the same receptors within the pituitary. Furthermore, we demonstrate that oral and nasal administration of GnRHa are viable alternatives to intracelomic injection for the stimulation of spermiation in the Fowler’s toad. These results not only serve to further inform our findings regarding hormone pathways but are promising for programs utilizing ART methods and present additional minimally-invasive options for hormone administration.

## 7. Author Contributions

Conceptualization, methodology, writing – original draft preparation, data curation, editing, and investigation, ARJ; Supervision, project administration, writing – editing, DK; Supervision, project administration, writing-review and editing, JMF; Supervision, project administration, writing-review and editing, STW; Resources, visualization, supervision, project administration, writing-review and editing, funding acquisition, AJK; Project conceptualization, resources, visualization, supervision, project administration, writing-review and editing, funding acquisition, CKK.

## 9. Acknowledgments

We would like to express our gratitude for the tireless efforts of our laboratory staff members for their assistance and animal care and the Mississippi State University School of Veterinary Sciences Pathology department for their assistance the histology component of this project. We would also like to thank Isabella Durham for her statistical modeling. Finally, we thank Dr. Seong Bin Park for his training and technical assistance in using the In-Vivo Imaging System (IVIS).

## 10. Data Availability Statement

Datasets are available on request.

## References

1. Al Shoyaib, A., Archie, S. R., Karamyan, V. T. (2020). Intraperitoneal Route of Drug Administration: Should it Be Used in Experimental Animal Studies? Pharm. Res., 37(1). 10.1007/s11095-019-2745-x

2. Anjum S, Krishna A, Tsutsui K. (2014). Inhibitory roles of the mammalian GnIH ortholog RFRP3 in testicular activities in adult mice. J Endocrinol. 223:79–91. doi:10.1530/JOE-14-0333

3. Browne, R. K., Seratt, J., Vance, C., Kouba, A. (2006). Hormonal priming, induction of ovulation and in-vitro fertilization of the endangered Wyoming toad (Bufo baxteri). Reprod. Biol. and Endocrinol., 4, 1–11. 10.1186/1477-7827-4-34

4. Browne, R. K. and Zippel, K. (2007). Reproduction and larval rearing of amphibians. ILAR J., 48(3), 214–234. 10.1093/ilar.48.3.214

5. Bronson, E., Guy, E.L., Murphy, K.J., Barrett, K., Kouba, A.J., Poole, V., Kouba, C.K. (2021). Influence of oviposition-inducing hormone on spawning and mortality in the endangered Panamanian golden frog (Atelopus zeteki). BMC Zool. 6, 17. 10.1186/s40850-021-00076-8

6. Burger, I. J., Chen, L. D., Lampert, S. S., Kouba, C. K., Barber, D., Smith, D., Cobos, C.,Kouba, A. J. (2023). Applying sperm collection and cryopreservation protocols developed in a model amphibian to three threatened anuran species targeted for biobanking management. Biol. Cons. 277. 109850. 10.1016/j.biocon.2022.109850

7. Burger, I., Julien, A. R., Kouba, A. J., Barber, D., Counsell, K. R., Pacheco, C., Krebs, J., Kouba, C. K. (2021). Linking in-situ and ex-situ populations of threatened amphibians through genome banking. Cons. Sci. & Prac. 3(11), 1–12. 10.1111/csp2.525

8. Campbell, L.G., Anderson, K. A., Marcec-Greaves, R. (2022). Topical application of hormone gonadotropin-releasing hormone (GnRH-A) stimulates reproduction in the endangered Texas blind salamander (Eurycea rathbuni). Cons. Sci. & Prac. 4(3), 1–8. 10.1111/csp2.609

9. Chen, D.M., Chen, L-D., Kouba, C.K., Songsasen, N., Roth, T.L., Allen, P.J., et al. (2024) Oral administration of GnRH via a cricket vehicle stimulates spermiation in tiger salamanders (Ambystoma tigrinum). PLoS ONE 19(7): e0289995. 10.1371/journal.pone.0289995

10. Clulow, J., Pomering, M., Herbert, D., Upton, R., Calatayud, N., Clulow, S., Mahony, MJ., Trudeau, VL. (2018). Differential success in obtaining gametes between male and female Australian temperate frogs by hormonal induction: a review. Gen. & Comp. Endocrinol. 265, 141–148.

11. Daniels, E., Licht, P. (1980). Effects of gonadotropin-releasing hormone on the levels of the plasma gonadotrophins (FSH) and LH) in the bullfrog, Rana catesbeiana. Gen. & Comp. Endocrinol. 42 (4), 455–463. 10.1016/0016-6480(80)90211-7

12. Della Togna, G., Howell, L. G., Clulow, J., Langhorne, C. J., Marcec-Greaves, R., Calatayud, N. E. (2020). Evaluating amphibian biobanking and reproduction for captive breeding programs according to the Amphibian Conservation Action Plan objectives. Theriogenol. 150, 412–431. 10.1016/j.theriogenology.2020.02.024

13. Gao Z, Zhang L, Sun Y. (2012). Nanotechnology applied to overcome tumor drug resistance. JCR.162(1):45–55. doi:10.1016/j.jconrel.2012.05.051

14. Graham, K.M., Langhorne, C.J., Vance, C.K., Willard, S.T., Kouba, A.J. (2018). Ultrasound imaging improves hormone therapy strategies for induction of ovulation and in vitro fertilization in the endangered dusky gopher frog (Lithobates sevosa), Cons. Physiol. 6(1), coy020, 10.1093/conphys/coy020

15. Helmer, P.J. and Whiteside, D.P. (2005). “Amphibian anatomy and physiology”. In: Clinical Anatomy and Physiology of Exotic Species, ed. B. O’Malley (Edinburgh : Elsevier Saunders), 3 – 14.

16. Homayun, B., Lin, X., Choi, H-J. (2019). Challenges and Recent Progress in Oral Drug Delivery Systems for Biopharmaceuticals. Pharmaceutics. 11(3):129. 10.3390/pharmaceutics11030129

17. Jin, J. F., Zhu, L. L., Chen, M., Xu, H. M., Wang, H. F., Feng, X. Q., Zhu, X. P., & Zhou, Q. (2015). The optimal choice of medication administration route regarding intravenous, intramuscular, and subcutaneous injection. Patient Prefer. Adherence. 9, 923–942. 10.2147/PPA.S87271

18. Julien, A. R., Kouba, A. J., Kabelik, D., Feugang, J. M., Willard, S. T., & Kouba, C. K. (2019). Nasal administration of gonadotropin releasing hormone (GnRH) elicits sperm production in fowler’s toads (Anaxyrus fowleri). BMC Zool. 4(1), 1–10. 10.1186/s40850-019-0040-2

19. Kouba, A. J., Vance, C. K., Willis, E. L. (2009). Artificial fertilization for amphibian conservation: Current knowledge and future considerations. Theriogenol. 71(1), 214–227. 10.1016/j.theriogenology.2008.09.055

20. Kouba, A.J., Vance, C.K., Calatayud, N.E., Rowlison, T.N.C., Langhorne, C. J., Willard, S. (2012). “Assisted reproduction technologies (ART) for amphibians” in Amphibian Husbandry Resource Guide, eds. V.A. Poole and S. Grow.

21. Kouba, A.J., Willis, E., Vance, C.K., Hasenstab, S., Reichling, S., Krebs, J. (2012). Development of assisted reproduction technologies for the endangered Mississippi gopher frog (Rana sevosa) and sperm transfer for in-vitro fertilization. RFD. 71, 214–227.

22. Lampert, S. S., Burger, I. J., Julien, A. R., Gillis, A. B., Kouba, A. J., Barber, D., Kouba, C. K. (2022). Sperm Cryopreservation as a Tool for Amphibian Conservation: Production of F2 Generation Offspring from Cryo-Produced F1 Progeny. MDPI Animals. 13(1), 53. 10.3390/ani13010053

23. Licht, P., Porter, D., Millar, R. P. (1987). Specificity of amphibian and reptilian pituitaries for various forms of gonadotropin-releasing hormones in vitro. Gen. & Comp. Endocrinol. 66(2), 248–255. 10.1016/0016-6480(87)90274-7F

24. Millar, R. P. (2003). GnRH II and type II GnRH receptors. Trends Endocrin. Met. 14(1), 35– 43. 10.1016/S1043-2760(02)00016-4

25. Millar, R. P., Lu, Z. L., Pawson, A. J., Flanagan, C. A., Morgan, K., Maudsley, S. R. (2004). Gonadotropin-releasing hormone receptors. Endocrin. Revs. 25(2), 235–275. 10.1210/er.2003-0002

26. Mitchell, M.A. (2009). Anesthetic Considerations for Amphibians. J. Exot. Pet Med., 18, 40–49.

27. Probst, C. E., Zrazhevskiy, P., Bagalkot, V., & Gao, X. (2013). Quantum dots as a platform for nanoparticle drug delivery vehicle design. Adv. Drug Deliv. Rev. 65(5), 703–18. doi:10.1016/j.addr.2012.09.036

28. Ramakrishnappa, N., Rajamahendran, R., Lin, Y. M., Leung, P. C. K. (2005). GnRH in non-hypothalamic reproductive tissues. Anim. Reprod. Sci. 88(1-2 SPEC. ISS.), 95–113. 10.1016/j.anireprosci.2005.05.009

29. Rowson, A. D., Obringer, A. R., Roth, T. L. (2001). Non-invasive treatments of luteinizing hormone-releasing hormone for inducing spermiation in American (Bufo americanus) and Gulf Coast (Bufo valliceps) toads. Zoo Biol. 20(2), 63–74. 10.1002/zoo.1007

30. Shaidani NI, McNamara S, Wlizla M, Horb ME. (2021). Obtaining Xenopus laevis Eggs. Cold Spring Harb Protoc. 1(3):pdb.prot106203. doi: 10.1101/pdb.prot106203. PMID: 33272976; PMCID: PMC7925371.

31. Silla, A. J., McFadden, M. S., Byrne, P. G. (2019). Hormone-induced sperm-release in the critically endangered Booroolong frog (Litoria booroolongensis): Effects of gonadotropin-releasing hormone and human chorionic gonadotropin. Cons. Physiol. 7(1), 1–10. 10.1093/conphys/coy080

32. Silla, A.J., Roberts, D.J., Byrne, P.G. (2020). The effect of injection and topical application of hCG and GnRH agonist to induce sperm-release in the roseate frog, Geocrinia rosea, Cons. Physiol. 8(1), coaa104, 10.1093/conphys/coaa104

33. Vieu, S., Le Poul, N., Tur, L. Aupée, C., Kerbrat-Copy, R., Bouhsina, N., Cojean, O., Fusellier, M. (2024). Ultrasound description of the coelomic cavity of the axolotl (Ambystoma mexicanum) in a clinically healthy population: a pilot study. Sci Rep. 14, 11787 10.1038/s41598-024-62264-z

34. Vu, M., & Trudeau, V. L. (2016). Neuroendocrine control of spawning in amphibians and its practical applications. Gen. and Comp. Endocrinol. 234, 28–39. 10.1016/j.ygcen.2016.03.024

35. Wright, K. N. and Whitaker, B. R. (2001). Amphibian Medicine and Captive Husbandry (Krieger Publishing Company).

36. Xing, Y., and Rao, J. (2008). Quantum dot bioconjugates for in vitro diagnostics & in vivo imaging. Canc. Biomark.: Sec. A of Dis. Markers 4 (6), 307–19.

